# Peanut Smut: A scientometric analysis for a pathosystem that concerns the Argentine peanut industry

**DOI:** 10.1101/2023.09.06.555881

**Authors:** Luis Ignacio Cazón, Juan Andrés Paredes, Esteban Miretti, Noelia Gonzalez, Lautaro Suarez, Cinthia Conforto, Alejandro Mario Rago

## Abstract

Since its first report in commercial batches in 1995, the prevalence and yield impact caused by smut disease have increased rapidly in peanut fields. At the same time, various working groups have studied this pathosystem using different approaches, contributing to the scientific knowledge of the disease. By recognizing the importance of a thorough bibliographic review and meticulous organization of information, the process of initiating new research projects becomes more effective. In light of this, the aim of this work was to provide a comprehensive scientometric analysis of the evolution of peanut smut research, spanning from its inception to the current day. For this purpose, we compiled bibliographic data about the disease and extracted information to calculate metrics. We observed that a smaller proportion of the scientific production was presented in peer-reviewed journals, the prevalent topics were epidemiology and breeding, and the collaborative endeavors were crucial for the scientific advancement in the study of this pathosystem. Additionally, the researchers with the most significant presence in the publications, the involved institutions, and the impact of the produced papers, among other trends were identified. Although there have been many scientific-technological advances in peanut smut over the years, this information is not reflected in scientific papers in peer-reviewed journals, which represents a great challenge for researchers involved in this topic. It is crucial to continue generating knowledge that contributes to the integrated management of this complex pathosystem. This will prevent further yield losses and the spread of the pathogen to new production areas.

## INTRODUCTION

Peanut smut, caused by *Thecaphora frezzii* Carranza & Lindquist, is considered the main sanitary concern affecting peanuts in Argentina. Its first report in commercial fields dates back to 1995, marking an unprecedented global event (Marinelli et al. 1995). Since then, its prevalence has gradually increased and it is currently recorded in all peanut fields within the peanut-growing areas of Córdoba province (Cazón et al. 2018, Paredes et al. 2022). The estimated yield losses caused by peanut smut can reach up to 35% in highly infested fields (∼50% smut incidence), and considering the peanut total production, the estimated losses exceed 14 million dollars (Paredes 2017). Since the peanut industry represents one of the most important regional economies in Argentina, the impact of this disease generated concern in the peanut community. In this context, different working groups began research projects considering different aspects of the disease to contribute to integrated management. During this process, much knowledge about the pathosystem was generated and published in different meetings and scientific journals. Typical approaches such as chemical control, peanut cultivar response, and disease epidemiology were initially considered, but contribution to short-term control was minimal (Marraro Acuña et al. 2007, 2009, Marinelli et al. 2008, Astiz Gassó et al. 2008, Oddino et al. 2008). This led the scientific community to deepen the studies that were being developed and to explore new areas, such as biology and physiology of the pathogen, biochemical and molecular aspects, and biological control, among other topics. At present, more than 25 years of dedicated research in this pathosystem have resulted in the development of various epidemiological and cultural tools, along with three registered smut-tolerant cultivars, a recommended fungicide, and numerous management recommendations for effective control of peanut smut.

The analysis of scientific production, especially in a complex subject like this, brings many advantages. Scientometric and bibliometric indicators provide researchers, development agencies, and institutions an objective insight into scientific advancements in a given field and their impact on practical applications (da Silva and Bianchi 2001, Glänzel 2003). Using this quantitative approach to scientific production, we can find out the number of scientific publications, the most studied topics, the institutions involved, and their interaction, among other metrics (Michás and Muñoz-Velasco 2013). Based on these indexes, it is possible to identify the areas where comprehensive studies need to be conducted, and allocate additional human resources or more financing as needed (Glänzel 2003). Considering the above-mentioned, and the importance of the systematization of knowledge, the objectives of this work were to carry out a scientometric analysis of peanut smut scientific production from the first report of the disease to the present.

## MATERIALS AND METHODS

A systematic review of peanut smut was carried out for this scientometric analysis. Only peer-reviewed articles and congress/conference proceedings were considered. Journalistic notes, fact sheets, or websites were excluded. The bibliographic search was carried out using Google (<u>https: /google.com.ar/</u>) and Google Scholar (<u>https: /scholar.google.com.ar/</u>). These search engines are widely used and recommended for this purpose, because of their ease of use, open access, and ability to analyze a wide scope of data (de Winter et al. 2014; Harzing and Alakangas 2016, Haddaway et al. 2015). The search was carried out using the keywords “peanut smut”, “carbón del maní” (Spanish translation), “carvão do amendoim” (Portuguese translation), “Thecaphora frezii” and “Thecaphora frezzii”. The publications found were archived in PDF format and the next information was extracted and systematized within a datasheet:

- Format of the publication: The publications were classified as congress/conference proceedings or peer-reviewed journal articles. The name of the congress/conference or journal was also considered to determine the most popular meeting/journal for the publication of peanut smut research results.
- General topic: To determine the most developed subjects in the scientific production of peanut smut, the title, full text, and keywords of each study were thoroughly examined. This information was also used to analyze the research fields of the most productive authors.
- Authors: For congress/conference publications, the first author was considered to analyze and rank the most productive researchers and institutions. For peer-reviewed articles, was also considered the total of authors in the paper, to analyze the contribution of each researcher to the scientific production.
- Impact of peer-review articles: The impact of a scientific article was studied through the number of citations (Moed 2005). This information was assessed by citation metrics analysis using Google Scholar on August 14^th^, 2023 (de Winter et al. 2014; Harzing and Alakangas 2016). In addition to the total number of citations, the citation factor was calculated (Del Ponte et al., 2017). This metric considers the number of citations per year since the article was published, thus eliminating time-related bias.
- Institutions involved and their collaborative interaction: Although there are many institutions involved, two approaches were analyzed: to determine the most productive institution, the affiliation of the first author was considered; to determine the interaction between institutions, the affiliation of each author in the publication was considered.

The analysis of the information and plots was carried out using the Rstudio software (R Core Team. 2021).

## RESULTS

Google Scholar returned 161 results from the bibliographic search. Out of the total, 25 (15.5%) were published in peer-reviewed scientific journals while 136 (84.5%) were congress/conference proceedings. The first publication was made in 1962, by Carranza & Lindquist in the Bulletin of the Argentine Botanical Society. This was the first report of the disease in wild peanuts worldwide. In 1995, approximately 33 years later, Marinelli et al. (1995) reported the disease in commercial peanuts for the first time. Subsequently, some epidemiological aspects of smut were documented in 2002 (Marinelli et al. 2002), but it was not until 2007 that a consistent pattern of yearly publication was observed (Fig. 1). According to Oddino et al. (2008), the prevalence of smut in the 2006/07 growing season, already identified the disease as a potential threat for peanut production in Córdoba. This may explain the interest in seeking tools and information that contribute to the integrated management of the disease. In consequence, the number of publications increased in 2010. In that year, ten abstracts on smut were presented at the XXIV National Peanut Conference in Argentina. This is a high number considering that only nine publications on the disease were produced until 2009 (one scientific article and eight abstracts in congresses/conferences) (Fig. 1). Considering the number of publications per year, 2021 was the most productive, with ten (out of 25) peer-reviewed articles and fourteen congress/conference proceedings published in that year (Fig. 1).

**Figure 1.**
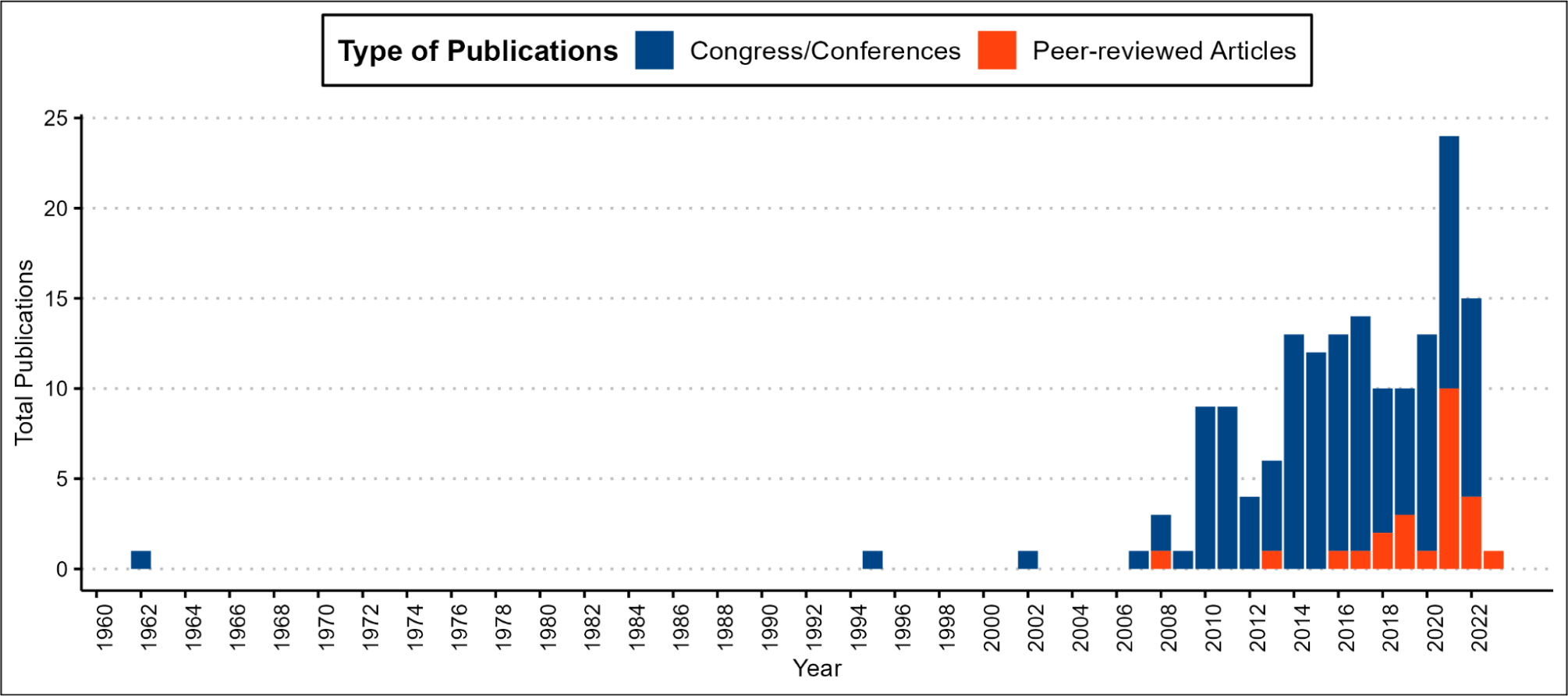
Frequency of publications on peanut smut from its first report to date. Different colors indicate different types of publications. Blue: Congress/conference proceedings. Red: articles in peer-reviewed journals.

### Addressed subjects

The general scientific production covered diverse topics (Fig. 2). The most studied subjects were epidemiology and breeding, which accounted for 38.5% (62/161) of the total publications. “Epidemiology” was the most developed topic (33/161) and had the highest number of publications over time, including 31 congresses/conference proceedings and two peer-reviewed articles from 2002 to the present. For “breeding”, scientific production began in 2009, but differently to the previous topic, more peer-reviewed articles were published (7/25). It is important to note that genetic resistance assumes an important role as an indispensable component in integrated disease management (Rago et al. 2017, De Blas et al. 2019, Kearney et al. 2021). Fortunately, Argentina has already registered three cultivars with high tolerance to peanut smut (Paredes 2022). Other important topics, such as chemical control and pathogen detection methods, have been systematically addressed over the years, but to a lesser extent. Biochemical aspects and biological control have gained importance in terms of the number of studies in recent years, with publications since 2014 and 2015 respectively (Fig. 3), aligning with the overall increase in scientific production.

**Figure 2.**
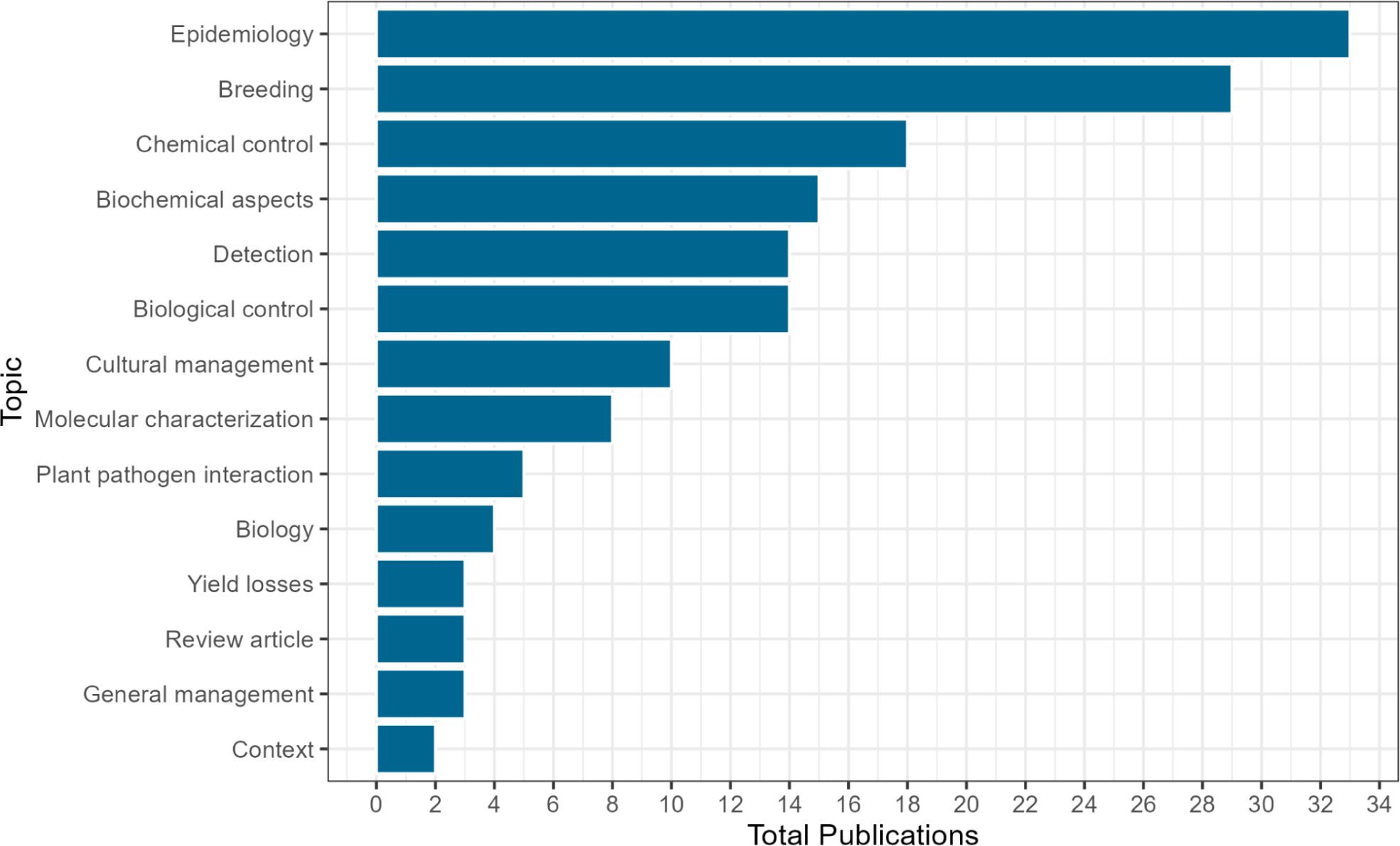
General subjects addressed in all publications (articles in peer-reviewed journals and proceedings at congresses/conferences) on peanut smut

**Figure 3.**
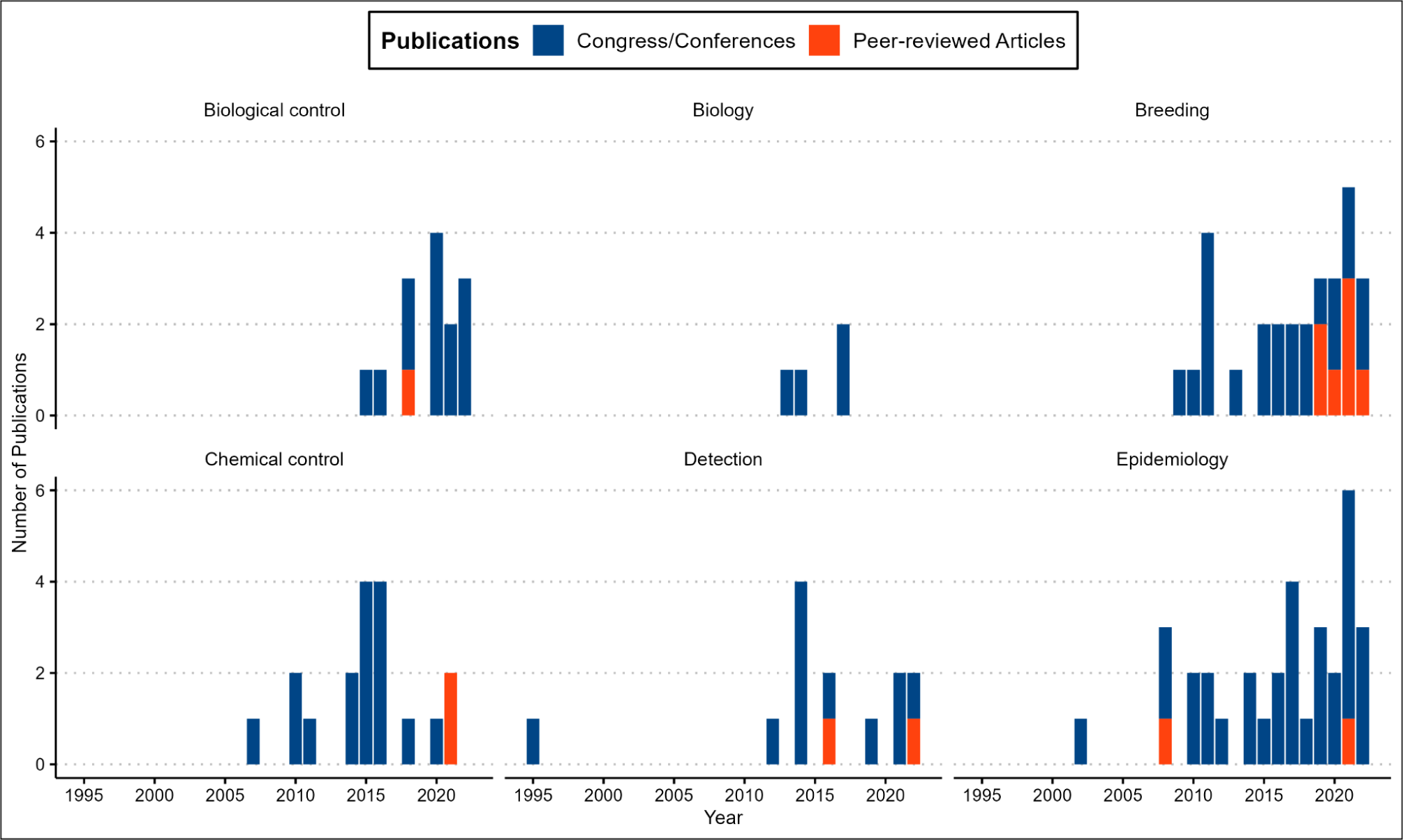
Frequency of publications of the six most addressed topics regarding peanut smut, sorted by year. Different colors indicate different types of publications. Blue: congress/conference proceedings. Red: articles in peer-reviewed journals.

### Conferences and congress proceedings

The first report of smut in cultivated peanuts date back to 1995 at the 7^th^ Congress of Mycology and the 17^th^ Argentine Conference on Mycology (Marinelli et al. 1995). Seven years later, the first epidemiological studies on the disease were published at the 17^th^ National Peanut Conference (Marinelli et al. 2002). This emerged as the predominant event for the presentation of results up to date, with 76.47% of the publications in this category (104/136). The second event, with a considerably lower number, is the Argentine Plant Pathology Congress, with 18.38% of the studies published (25/136). The remaining 5.15% of the publications (6/136) were attributed to other events (S1).

To explore the most productive researchers in this category, the first author of each publication was considered. Paredes JA (INTA-IPAVE) is in first place with 21 publications between 2014-2022 (2.3 publications per year). The main topics addressed by this author were epidemiology and chemical control. The second most productive author is Marraro Acuña F (INTA-Manfredi), with 10 publications between 2007 and 2014 (1.4 publications per year) covering various topics, such as breeding, epidemiology, plant-pathogen interaction, and chemical and cultural control. In third place, with 8 studies published as first authors are Oddino C (UNRC), Mary V (UNC), Cazón LI (INTA-IPAVE), and Astiz Gasso (UNLP) (Fig. 4) (S1). A total of 136 publications in this category involving 53 first authors, 62 out of the 136 studies (45.5%) are authored by the six researchers mentioned above, indicating their commitment and consistency in advancing the understanding of this pathosystem over time.

**Figure 4.**
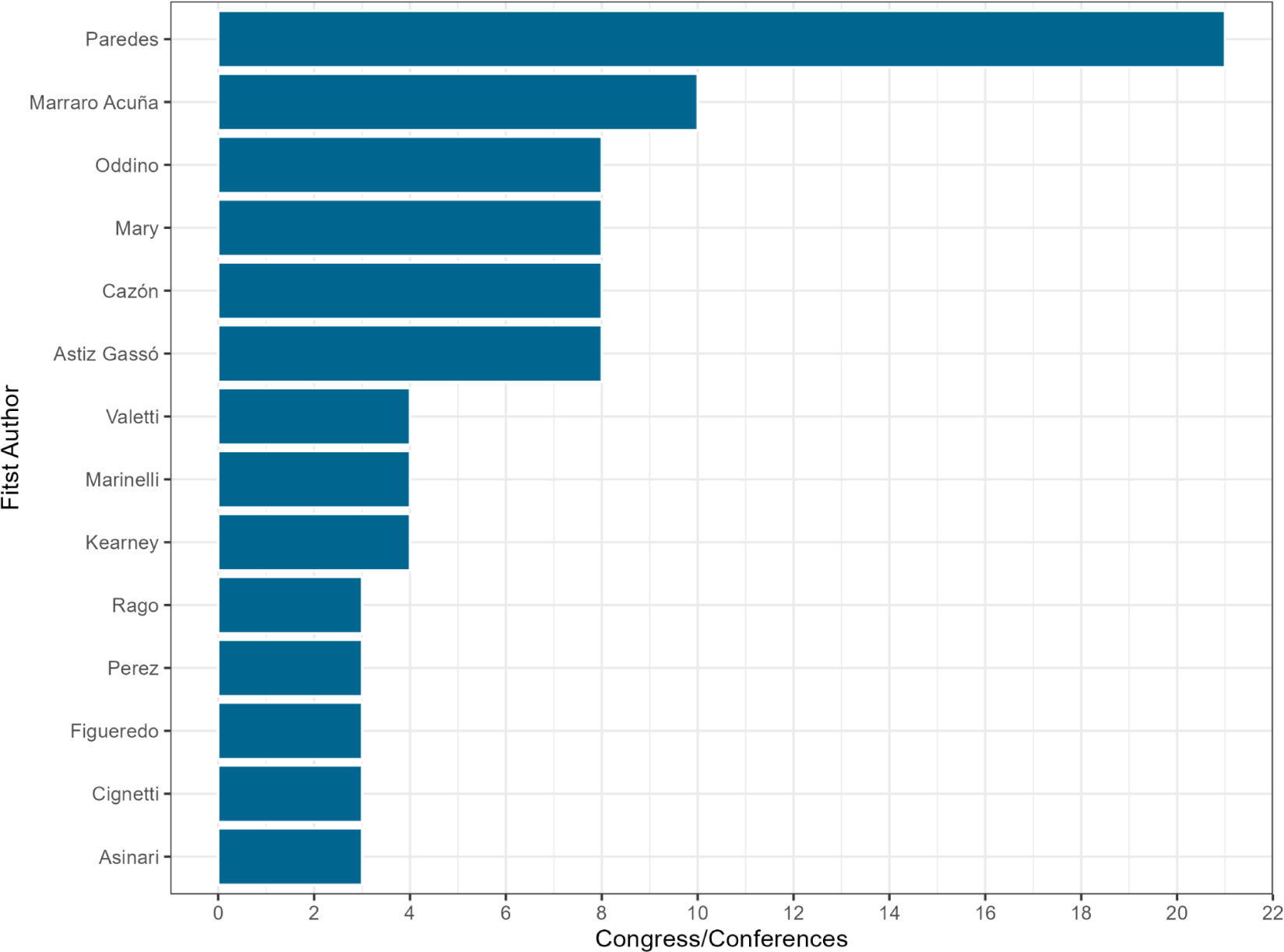
List of first authors with at least three publications in congress/conferences.

### Peer-reviewed journal articles

In this category, 25 articles published in peer-reviewed journals were found. Most of them were published in Peanut Science (4), the European Journal of Plant Pathology (3), and Plant Disease (2). The rest of the publications were in other journals listed in S2. All articles, except the first in Spanish by Marinelli et al. (2008) in AgriScientia journal, were published in English. The next publication in this category was made in 2013 by Conforto et al. (2013). From 2016, at least one article per year has been published up to the present. (Fig. 1).

Considering the first author of the publications, the researchers with the highest production are Soria N (UCC) with three published papers. Paredes JA (INTA-IPAVE), De Blas F (UNC), Arias R (USDA), and Cazón LI (INTA-IPAVE) have each contributed two articles. The rest of the authors are listed in S2. Since the production of papers is low and it is not possible to observe a significant difference in scientific production considering the first authors, we decided to analyze the contribution of each researcher in the overall production of articles, without considering the order in the list of authors. Using this approach, we counted 86 researchers present in the 25 papers found. Considering the top three, Rago AM (INTA-CIAP) was the one with the greatest presence, whose name was included in ten published papers, followed by Oddino C (UNRC) with presence in eight articles. In the third position, Cazón LI and Paredes JA (INTA-IPAVE), are presents in seven papers (Fig. 5).

**Figure 5.**
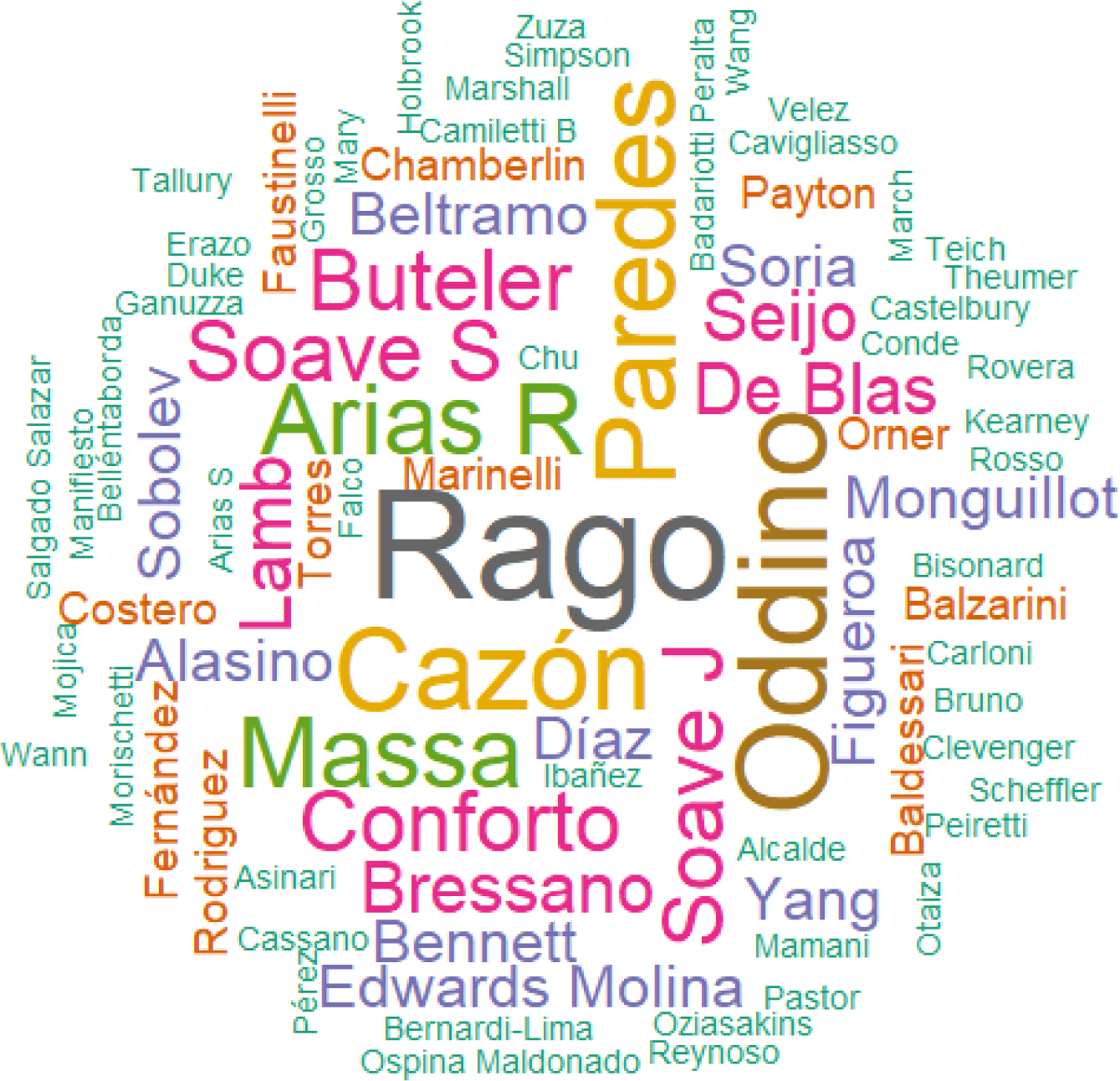
Word cloud represents the contribution of each researcher to the production of peer-reviewed scientific articles. The size of the names represents the contribution level, with larger names indicating greater contributions. Names with the same color signify equal contributions.

### Impact of peer-review articles

The articles with the greatest impact can be seen in (Fig. 6). Considering the total number of citations, Rago et al. (2017), Conforto et al. (2013) and Marinelli et al. (2008) are the most cited articles, with 34, 22, and 20 citations respectively (Fig 6A). In terms of the citation factor (number of citations per year), Rago et al. (2017) continue in the first place, with a factor of 5.7 citations per year, followed by Paredes et al. (2021), Bressano et al. (2019), and De Blas et al. 2021, with citation factors of 4.5, 4.0 and 4.0 citations per year respectively (Fig. 6B).

**Figure 6.**
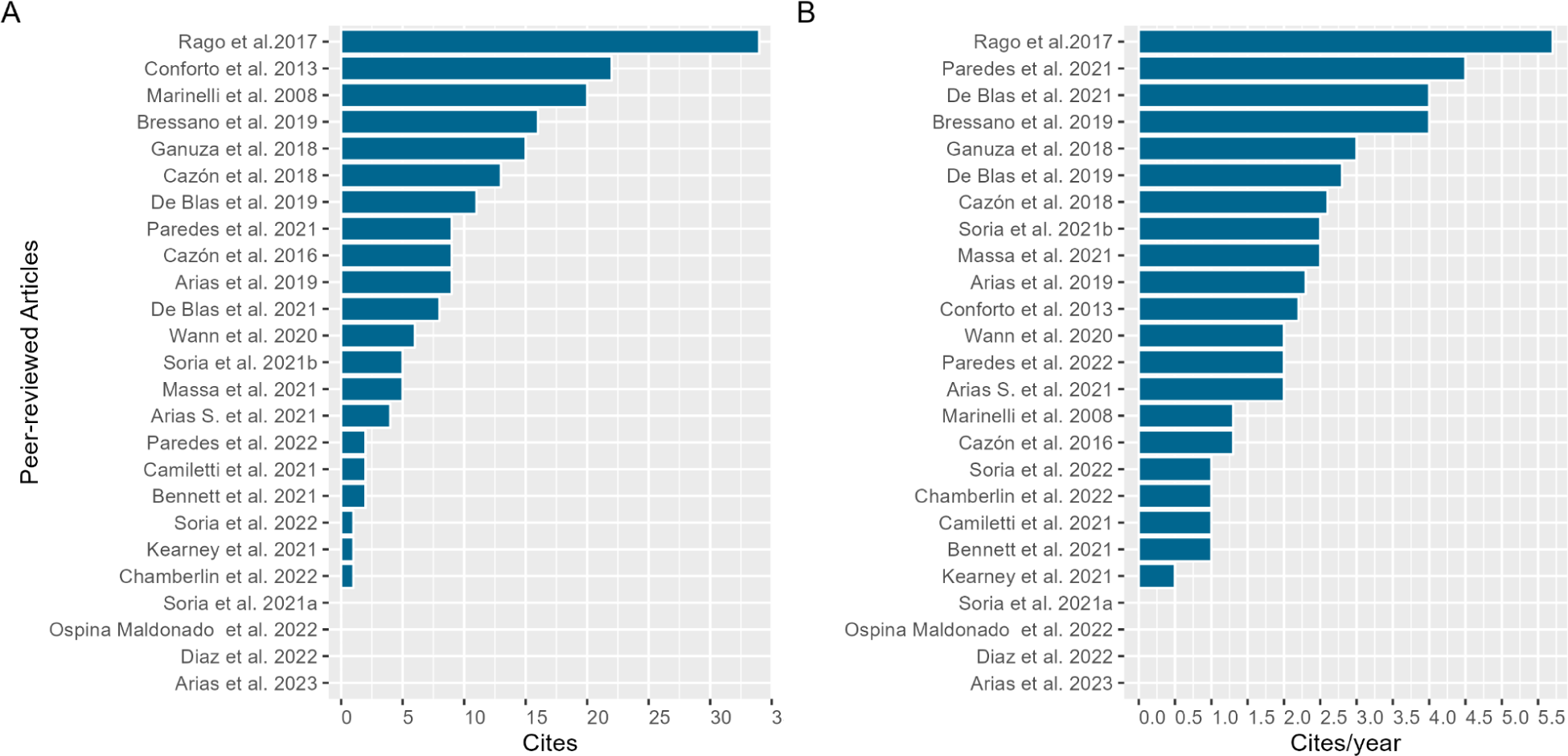
Total number of citations to scientific papers since their publication (A) and number of citations per year for each article (B).

### Institutions involved in scientific production and their interaction

Considering the first authors in congress/conferences and articles in peer-reviewed journals, researchers affiliated with INTA were the most prolific (Fig. 7). Of the 25 peer-reviewed articles found, six correspond to INTA-CIAP, five to USDA and Universidad Nacional de Córdoba (UNC). Universidad Nacional de Río Cuarto (UNRC) and Universidad Católica de Córdoba (UCC) each institution published three papers. Finally, CEPROCOR, Universidad Simón Bolívar, and International Peanut Growers (IPG) have each contributed a paper (Fig. 7A). Of the 136 publications in congress/conferences, the first authors with INTA affiliation presented 65 works (53 to CIAP and twelve to EEA-Manfredi), followed by UNRC (30), UNC (19), Universidad Nacional de La Plata (UNLP) (9), Criadero El Carmen (6), CEPROCOR (4), Universidad Nacional de Villa María (3), AGD (3), USDA (1), UCC (1), OLEGA (1) and Instituto de Botánica del Nordeste (IBONE) (1) (Fig. 7B).

**Figure 7.**
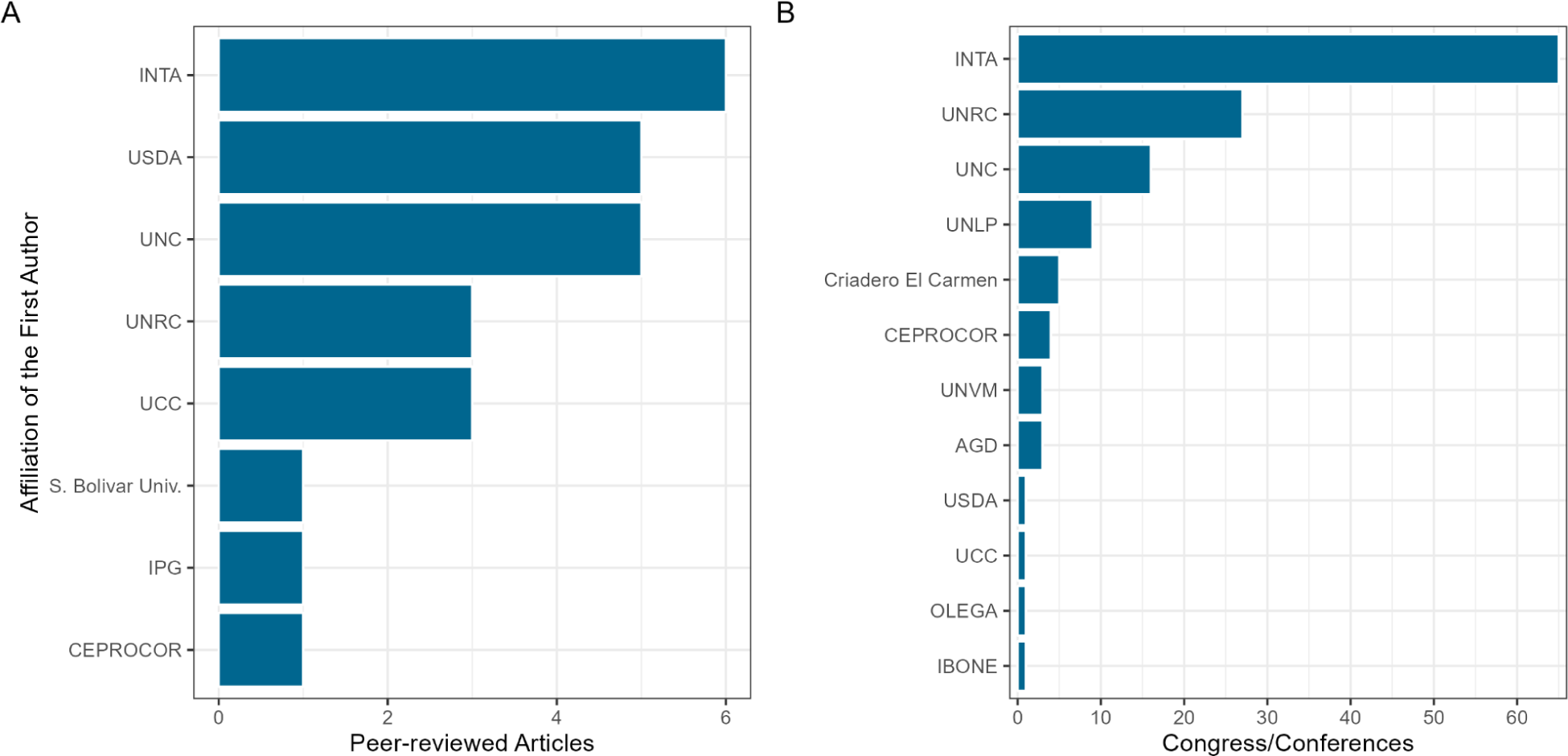
Contribution of each institution to the production of peer-reviewed scientific articles (A) and congress/conference proceedings (B).

As previously mentioned, there are many institutions working synergistically to generate knowledge concerning this pathosystem of knowledge about this pathosystem. Analyzing the interaction between institutions (considering the filiation of each author listed on publications) we observed that in Argentina, 68.32% of the publications (101 congresses/conferences and nine peer-reviewed articles) arose from the interaction between public agencies, while 19.25% (27 congresses/conferences and four peer-reviewed articles) have emerged from public-private interaction. In terms of international cooperation (ARG-USA), 6.21% of the studies result from this type of interaction. The difference is that this percentage primarily comprises scientific articles published in peer-reviewed journals (one abstract in congress/conference and nine peer-reviewed articles). Two peer-reviewed articles and seven congresses/conference proceedings (5.59%) were carried out without cooperation between institutions since all the authors had the same affiliation. Finally, only one article (review) was published in association between institutions from the USA and Colombia in 2022 (Fig. 8).

**Figure 8.**
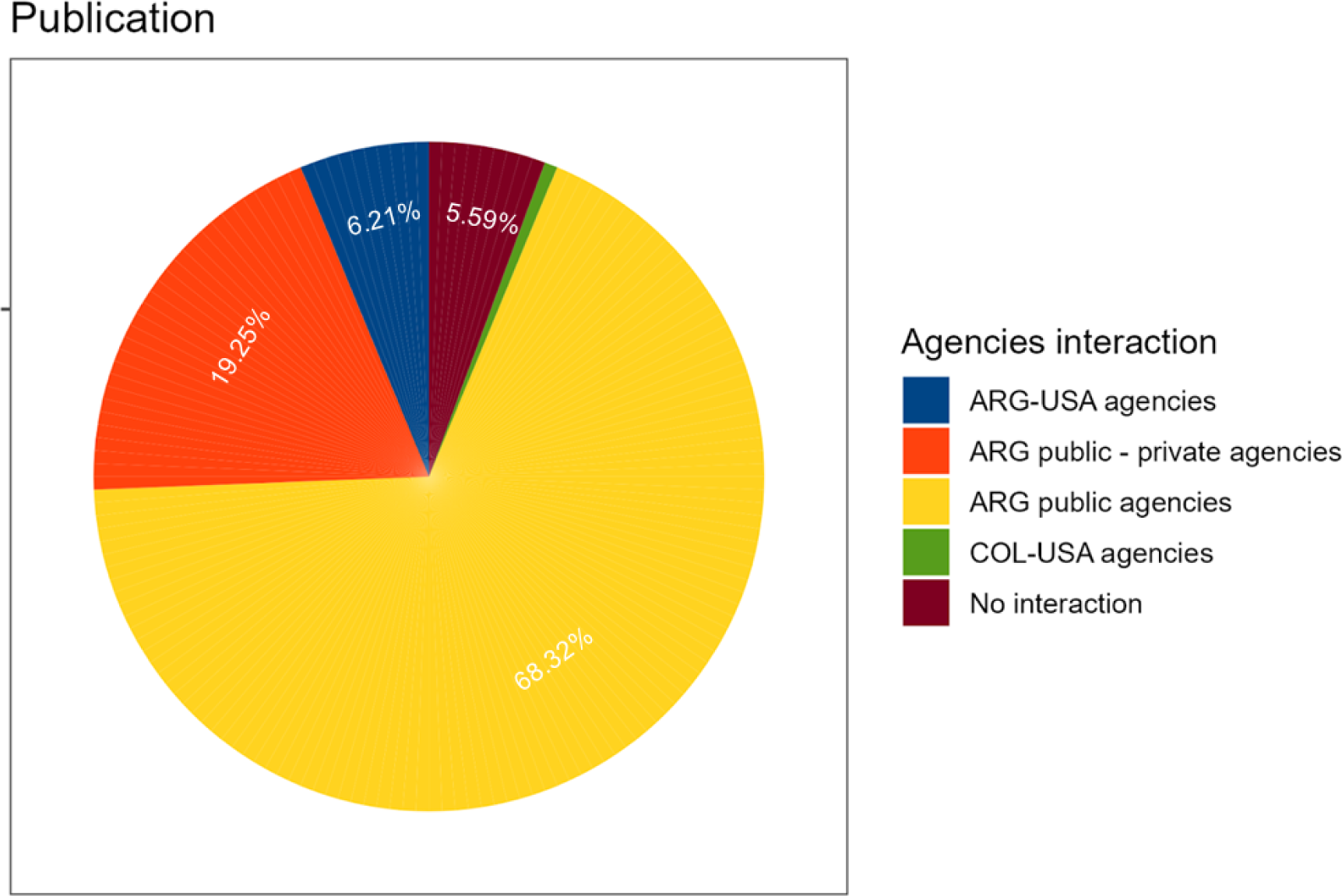
Cooperation among institutions in scientific production

## DISCUSSION

In this work, many important scientometric and bibliometric variables in relation to the scientific production of peanut smut were determined. Considering the importance of peanuts as one of the main regional economies of Argentina, this information and its analysis are crucial, mainly for two reasons. First, Argentina is the only country that has reported the disease in commercial peanuts (Cazón et al. 2018). Although the fungus was observed in Brazil and Bolivia, the disease has only been reported in wild peanuts (Carranza and Lindquist, 1962; Soave et al. 2014). The second reason is related to the export characteristics of the industry in this country.. According to data from Calzada and Rozadilla (2018), more than 90% of Argentine peanut production is used for this commercial activity. Although in recent years a policy promoting local consumption has been observed, a large part of the production continues to be destined for the international market (Camara Argentina del maní 2022). The convergence of these two factors drives Argentina to assume a leading role in exploring this pathosystem. In fact, it is Argentine researchers and institutions that predominantly drive scientific advancements in the field of peanut smut. Although few studies were produced by international institutions, the most part included Argentine researchers as co-authors. (Arias et al. 2019, 2023, Bennett et al. 2021, Chamberlain et al. 2022, Massa et al. 2021, Wan et al. 2020).

During the bibliographic compilation process, we were able to observe that the number of publications in peer-reviewed journals is low. After 28 years, just 25 articles have been published on the disease (less than one publication per year). Only starting from 2016, up to the present, the publication of at least one article per year is noticeable. In fact, over 90% of the published articles on smut (23 out of 25) were presented in this period (last 8 years). Regarding the production peak observed in 2021, not only for peer-reviewed articles but also in congress/conference proceedings, it is important to mention that during that year (and the previous one) the health policy adopted by Argentina against the COVID-19 pandemic recommended home office activities. This led many researchers to focus on data analysis and publication of the results obtained up to that moment.

In 2017, we observed that two significant events had a direct influence on the acceleration of scientific production from that year. The first was the hosting of the Ninth International Conference of the Peanut Research Community in Córdoba, Argentina. Here, the problem of peanut smut was presented for the first time to the international scientific community. In addition to sharing ongoing research, the conference also facilitated field visits, providing attending researchers with firsthand exposure to severely affected plots, which left a strong impact on the participants. It was during this event that a substantial portion of the connections with the USDA were formalized, leading to the publication of 9 articles starting in 2019. The second important event was the publication of the review conducted by Rago et al. (2017). This article exposed, for the first time, the economic impact of peanut smut on the industry and the complexity of the pathosystem in terms of disease management. Local advancements in diverse fields including pathogen biology, molecular aspects, epidemiology, integrated management, and chemical control, among other topics, were also published. The importance of this publication for subsequent scientific development can be observed through the high number of citations and the citations/year factor (Fig. 6).

Regarding congress/conference proceedings, we can observe that works falling within this category significantly outnumber publications in peer-reviewed journals. Generally, these encompass progress reports of ongoing research, which often culminate in peer-reviewed journal publications. As indicated by Moed et al. (2005), the incorporation of congress/conference proceedings in the scientometric analysis allows us to identify trends, new lines of development, and opportunities to foster synergy through collaborative spaces focused on specific subjects. An average of 10 abstracts per year were published since 2010, showing a growing trend up to the present. Out of the 129 congress/conference proceedings in this period, 104 (80.6%) were given at the Jornada Nacional del Maní (Argentina), underscoring the significance of this event for presenting preliminary results. While the upward publication trend in this category continues, a decline in presentations related to chemical control was observed starting in 2019. This is likely due to the release, in 2018, of the only commercial fungicide registered for controlling peanut smut. Concurrently, an increase in works focusing on biological control as a developing alternative for disease management has been seen (Fig. 9). This was not the case for breeding, as the first smut-tolerant cultivar was released in the same year, however, the number of works presented at congress/conferences remained constant up to the present. Even in light of the availability of new cultivars and advances in breeding programs, it remains crucial to persist in the development of other areas. Pathogen characteristics related to inoculum generation, its accumulation in fields, and spore dispersion dynamics contribute to a change in the epidemiological context over time. Therefore, incorporating management tools would help mitigate the disease impact in new scenarios, especially in new production areas.

**Figure 9.**
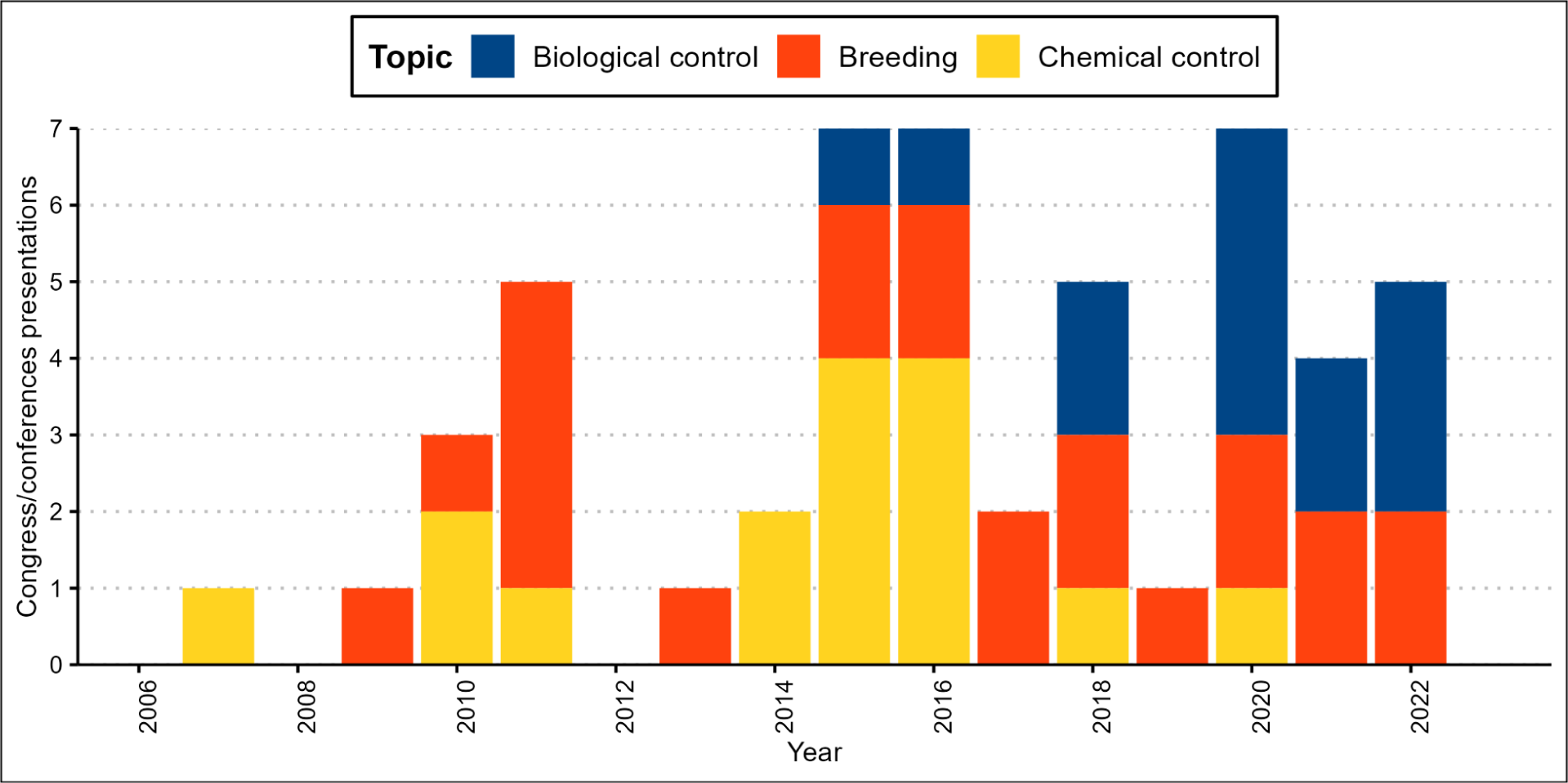
Frequency of congress/conference proceedings addressing Biological control (blue bars), Breeding (red bars), and Chemical control (yellow bars).

In a broader context, from its first report on commercial peanuts in 1995 until 2010, only a few studies were conducted on this pathosystem. Being a “new disease”, the initial objective was to comprehend its behavior in the field and certain patterns related to the infection processes (Astíz Gassó et al. 2008, Marinelli et al. 2008, 2002, 1995, Marraro Acuña et al. 2009, 2007, Oddino et al. 2008). This is why the initial studies primarily focused on epidemiological aspects, which served as a basis to demonstrate the potential impact of the disease on the crop and the need to allocate more resources for its research. Another area explored from the outset was breeding. The significance of tools generated in this discipline is evident in the number of publications in peer-reviewed journals compared to the total publications in this category (Chamberlin et al. 2022, De Blas et al. 2021, 2019, Kearney et al. 2021, Massa et al. 2021, Wann et al. 2020, Bressano et al. 2019). As previously mentioned, Argentina boasts three registered cultivars with significant tolerance to peanut smut, which is essential for effective disease management. Nevertheless, to harness their benefits and promote the sustainability of tolerance, the use of these cultivars must occur within an appropriate framework of integrated management.

The third most developed area is chemical control (Fig. 2). Despite this, the majority of the publications are found in the category of congress/conference proceedings. Only two articles have been published in a peer-reviewed journal studying application technology and control efficacy of different chemical groups (Camiletti et al. 2021, Paredes et al. 2021). Regarding the abstracts presented at congress/conferences, a strong emphasis was placed on the application technology, including types of nozzles, the timing of application, and formulation, in addition to the control efficacy of the products. One explanation for the low production of scientific articles on this topic might lie in the variability of results offered by chemical fungicides under different conditions of humidity, initial inoculum, timing, and method of application, among other factors (Paredes 2022), which hinders the formulation of consistent conclusions. As mentioned earlier, there is only one recommended fungicide for peanut smut (triadimenol 30% + myclobutanil 20%). It was released in 2018 when the epidemiological context had few highly infested fields. Currently, the polyetic characteristic of the pathogen has altered the situation, reinforcing the need for new management tools.

In terms of the impact of the published works, the one produced by Rago et al. (2017) holds the highest number of total citations and the highest citations/year factor. This underscores the aforementioned importance of this publication in subsequent scientific production processes related to the pathogen, especially in peer-reviewed journals. This can be observed in the bibliographic references of 17 out of the 20 papers produced on the disease from 2018 onwards, where Rago et al. (2017) are included. Works addressing chemical control (Paredes et al. 2021a) and breeding (Bressano et al. 2019, De Blas et al. 2021) follow in the rankings of the citations/year factor. As previously mentioned, these topics rank among the most frequently studied in scientific research on peanut smut, indicating their considerable significance.

As we know, there are numerous institutions involved in the pursuit of knowledge regarding this pathosystem. While authors affiliated with INTA lead in the first authorships of peer-reviewed articles and congress/conference proceedings (Fig. 7AB), collaborative processes have been crucial in the scientific development of this pathosystem. Our results show that most of the scientific production was carried out through collaboration between public institutions in Argentina, but only nine articles were submitted to peer-reviewed journals. Different situations exist regarding public-private and international cooperation. Despite low interaction in scientific production, four and eight articles were respectively published under peer review. This demonstrates the importance of combining skills and approaches in research processes. These existing connections, their consolidation, and the creation of new links are essential in the search for solutions to the peanut smut problem. The complexity of disease management requires an interdisciplinary approach to define effective strategies that reduce their intensity in the fields and address concerns such as contamination of new production areas (Paredes 2022).

Although many topics still require more in-depth study, significant progress has been made in understanding this pathosystem. Currently, one of the main challenges is to encourage various research groups to systematize and publish their results (with appropriate methodological rigor) in peer-reviewed scientific articles format. Consequently, all the efforts invested in peanut smut research thus far, along with ongoing work, would become significant not only at the local level but also within the international scientific community. This would bring forth numerous benefits to the peanut industry and scientific advancement.

## Author’s contribution

LIC conceptualized the work, conducted the bibliographic research, analyzed the data and wrote the manuscript; JAP contributed to data analysis and wrote the manuscript; EM, NG, and LS contributed to the bibliographic search and analysis of each publication; CC and AMR conceptualized the work and wrote the manuscript.

## Acknowledgments

We wish to thank INTA and Fundación Maní Argentino for providing resources for compiling this project, and Boris Camiletti, for critically reviewing the manuscript prior to submission.

## Supplementary information

**S 1.**
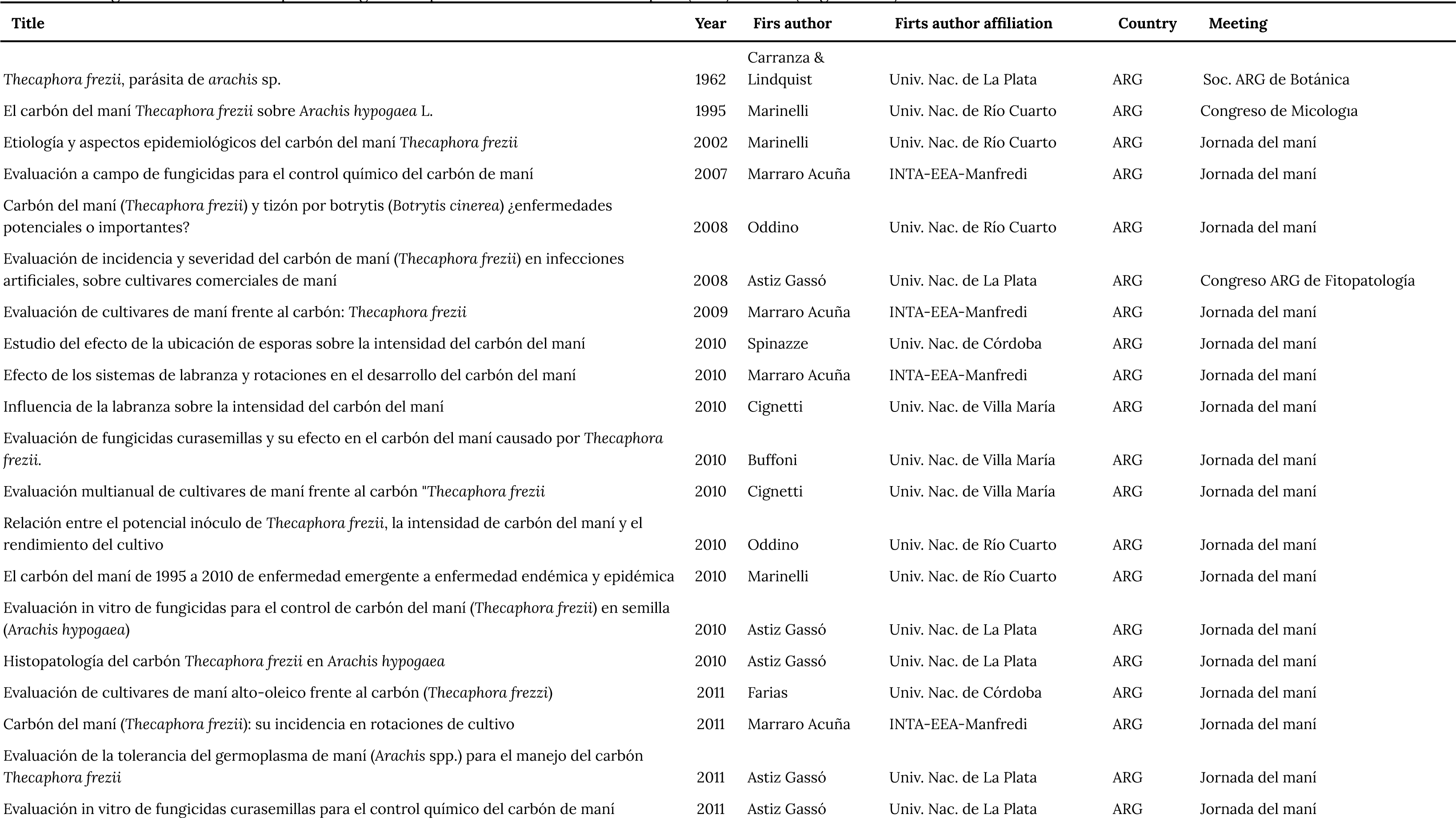
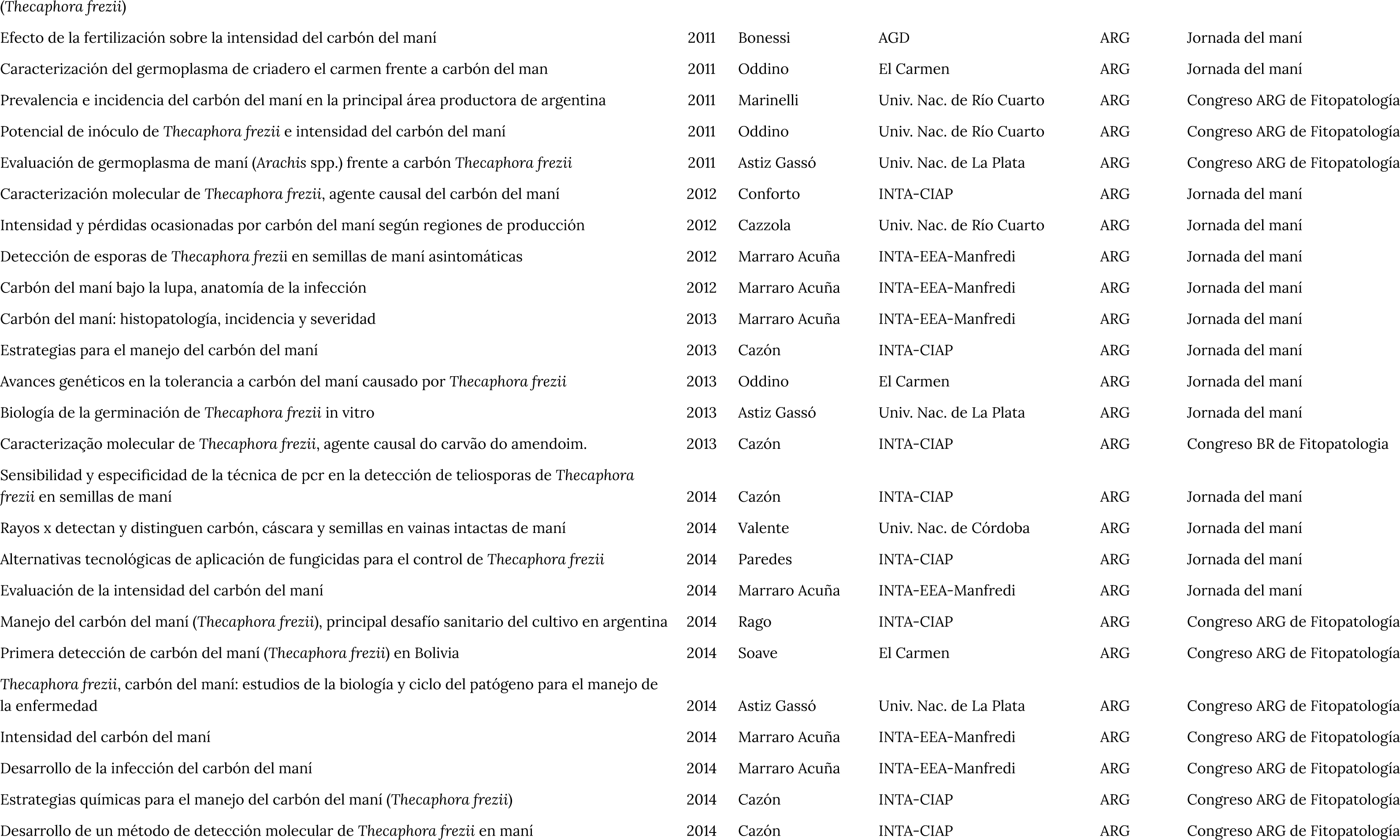
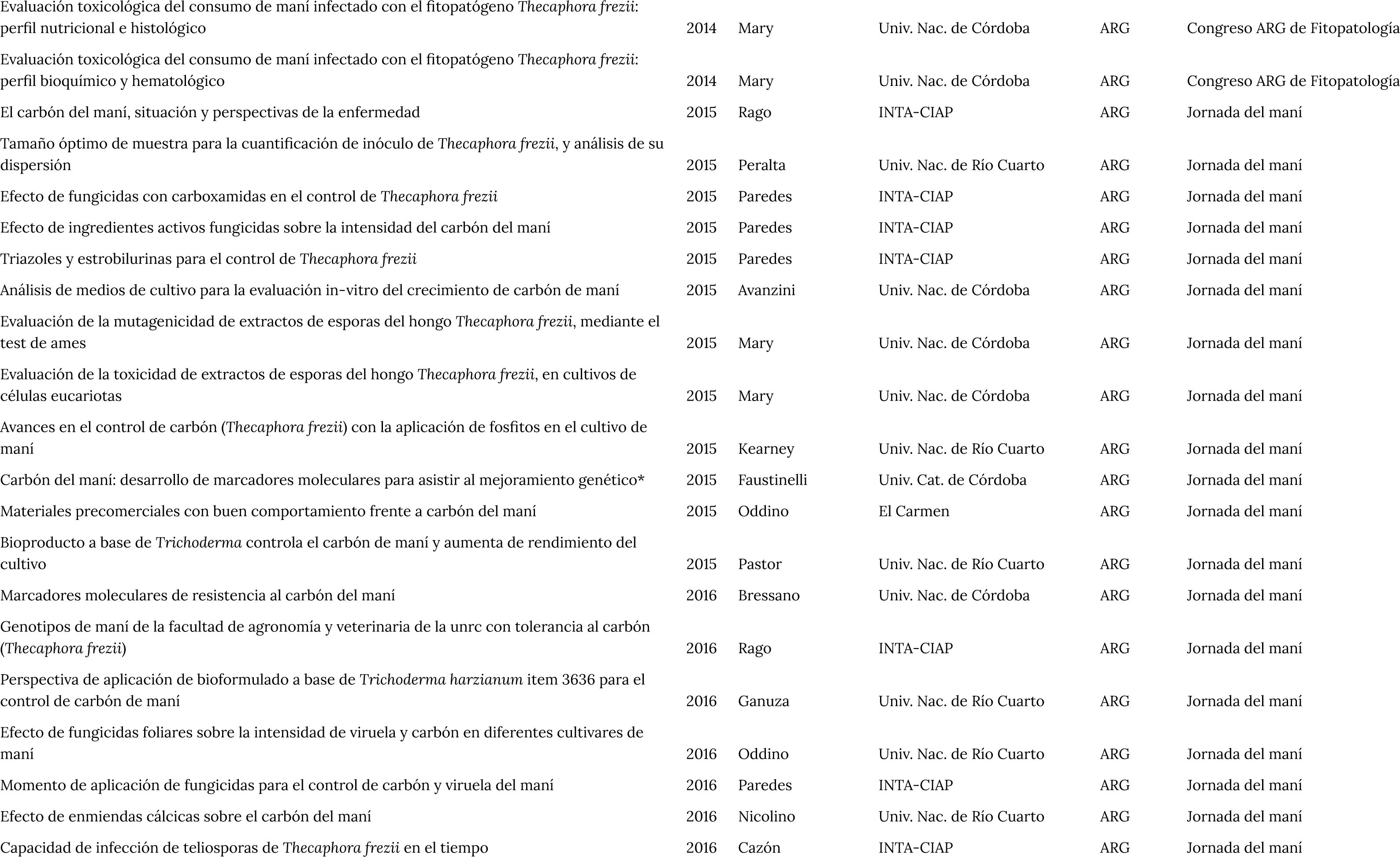
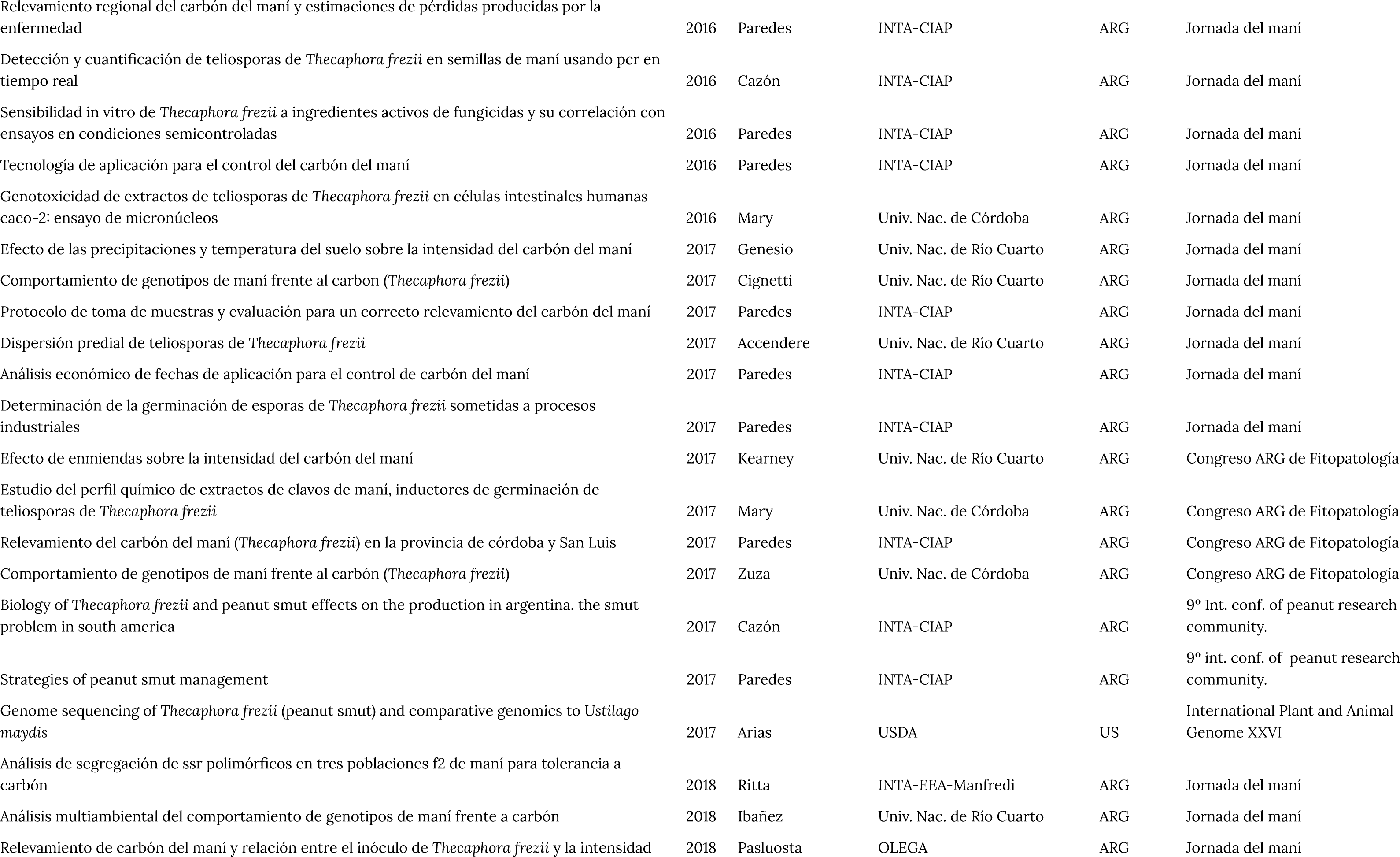
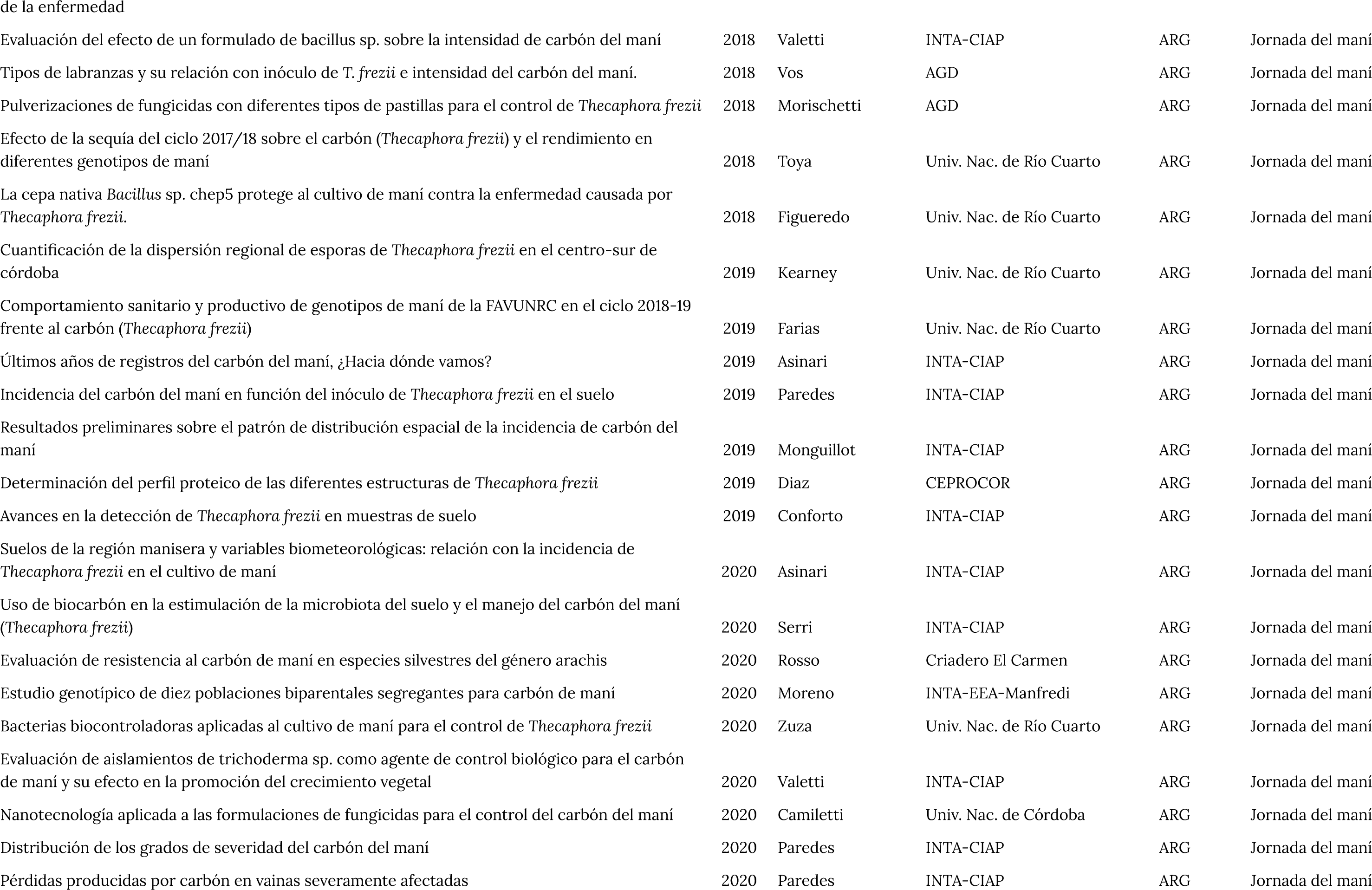
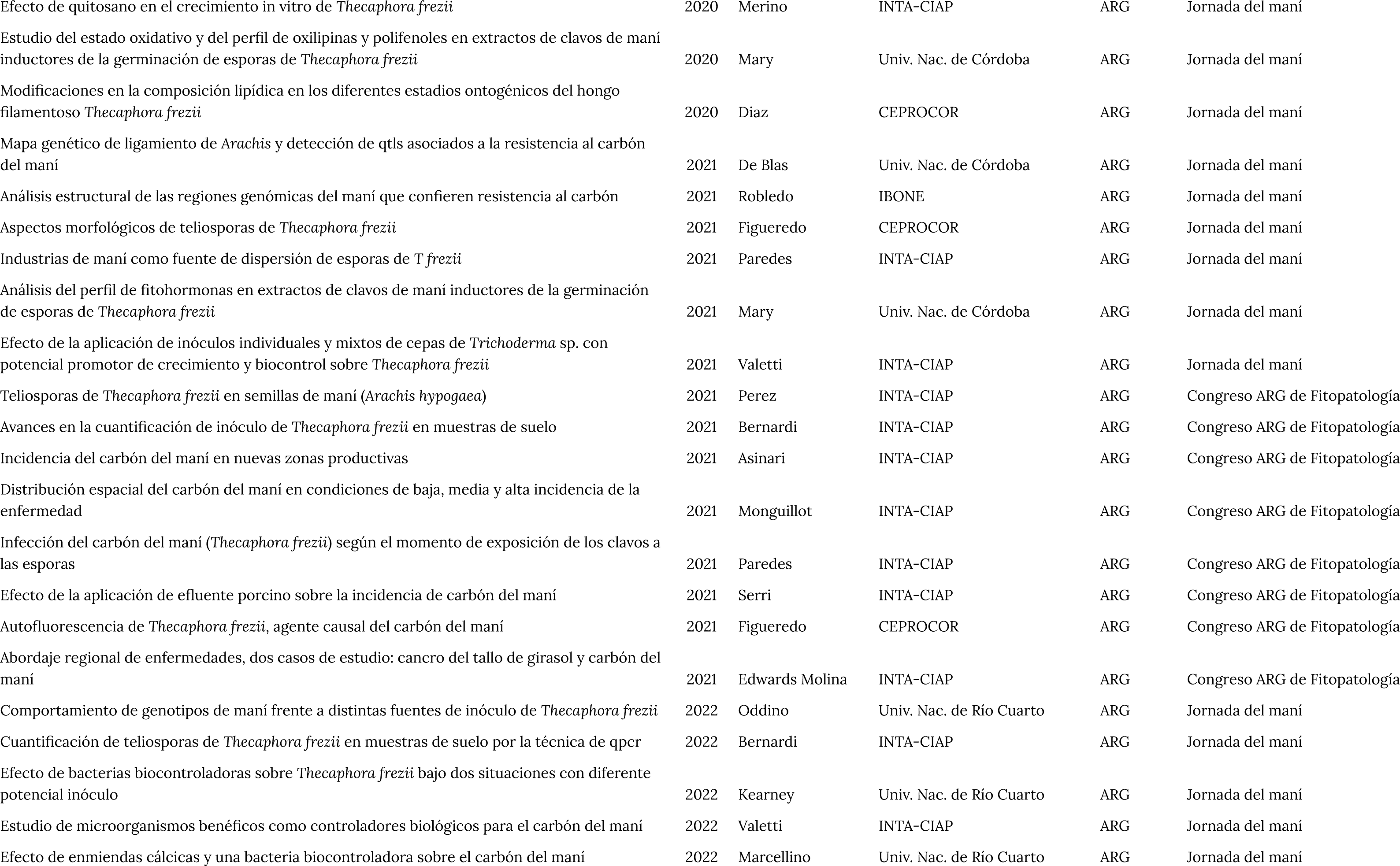
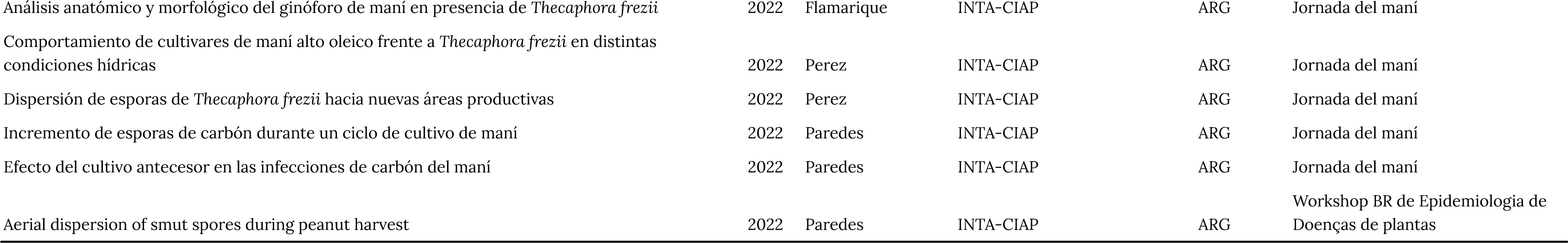
List of congress and conference proceedings about peanut smut from its first report (1962) to date (August 2023).

**S 2.** List of peer-reviewed articles published about peanut smut.

S 2. List of peer-reviewed articles published about peanut smut.
| Title | Year | First author | First author affiliation | Country | Journal | Total n° of cites | Factor cites/year |
| --- | --- | --- | --- | --- | --- | --- | --- |
| Aspectos biológicos y epidemiológicos del carbón del maní (*Arachis hypogaea* L.) causado por *Thecaphora frezii* Carranza & Lindquist | 2008 | Marinelli | Univ. Nac. de Río Cuarto | ARG | Agriscientia | 20 | 1.3 |
| Molecular sequence data of *Thecaphora frezii* affecting peanut crops in argentina | 2013 | Conforto | INTA-CIAP | ARG | European J of Plant Path | 22 | 2.2 |
| Molecular detection of *Thecaphora frezii* in peanut (*Arachis hypogaea* L.) seeds | 2016 | Cazón | INTA-CIAP | ARG | J of Plant Path | 9 | 1.3 |
| Peanut smut : from an emerging disease to an actual threat to argentine peanut production | 2017 | Rago | INTA-CIAP | ARG | Plant Disease | 34 | 5.7 |
| The biology of *Thecaphora frezii* smut and its effects on Argentine peanut production | 2018 | Cazón | INTA-CIAP | ARG | Intech Open | 13 | 2.6 |
| Efficacy of the biocontrol agent *Trichoderma harzianum* item 3636 against peanut smut, an emergent disease caused by *Thecaphora frezii* | 2018 | Ganuza | Univ. Nac. de Río Cuarto | ARG | European J of Plant Path | 15 | 3.0 |
| Mitogenome and nuclear-encoded fungicide-target genes of *Thecaphora frezii* causal agent of peanut smut | 2019 | Arias | USDA | US | Fungal Genomic and Biol | 9 | 2.3 |
| Identification of smut resistance in wild arachis species and its introgression into peanut | 2019 | De Blas | Univ. Nac. de Córdoba | ARG | Crop Science | 11 | 2.8 |
| Introgression of peanut smut resistance from landraces to elite peanut cultivars (*Arachis hypogaea* L.) | 2019 | Bressano | Univ. Nac. de Córdoba | ARG | Plos One | 16 | 4.0 |
| Phenotypic variation of peanut smut (*Thecaphora frezii*) incidence and severity in the u.s. peanut mini-core collection | 2020 | Wann | International peanut growers | US | Peanut Science | 6 | 2.0 |
| Where does the peanut smut pathogen *Thecaphora frezii* fit in the spectrum of smut diseases | 2021 | Arias S | Univ. Nac. de Córdoba | ARG | Plant Disease | 4 | 2.0 |
| A note on the association between *Thecaphora frezii* infection and peanut | 2021 | Bennett | USDA | US | PeanScie | 2 | 1.0 |
| Fungicide efficacy of nanocrystal-based formulations against peanut smut | 2021 | Camiletti | UNC | ARG | Crop Protection | 2 | 1.0 |
| Genetic mapping and QTL analysis for peanut smut resistance | 2021 | De Blas | Univ. Nac. de Córdoba | ARG | BCM Plant Biology | 8 | 4.0 |
| Response of peanut (*Arachis hypogaea* L.) genotypes to smut (*Thecaphora frezii*) in the peanut growing region of argentina | 2021 | Kearney | Univ. Nac. de Río Cuarto | ARG | Peanut Science | 1 | 0.5 |
| Genotyping tools and resources to assess peanut germplasm: smut-resistant landraces as a case study | 2021 | Massa | USDA | US | Peer J | 5 | 2.5 |
| Efficacy of fungicides against peanut smut in Argentina | 2021 | Paredes | INTA-CIAP | ARG | Crop Protection | 9 | 4.5 |
| Relationship between incidence and severity of peanut smut and its regional distribution in the main growing region of argentina | 2022 | Paredes | INTA-CIAP | ARG | Tropical Plant Path | 2 | 1.0 |
| Identification and expression of the fungal central effector pep1 in *Thecaphora frezii* | 2021a | Soria | Univ. Cat. de Córdoba | ARG | Method | 0 | 0 |
| Identification of chitin synthase and chitinase genes in three ontogenetic stages from *Thecaphora frezii*, the causal agent of peanut smut disease | 2021b | Soria | Univ. Cat. de Córdoba | ARG | Physiol and Molec Plant Path | 5 | 2.5 |
| Identification of germplasm resistant to peanut smut | 2022 | Chamberlin | USDA | US | Peanut Science | 1 | 1.0 |
| Peanut smut: a diagnostic guide | 2022 | Ospina | Univ. Simón Bolívar | COL | Plant Health Progress | 0 | 0 |
| Elucidating potential effectors, pathogenicity and virulence factors expressed by the phytopathogenic fungus *Thecaphora frezii* through analysis of its transcriptome | 2022 | Soria | Univ. Cat. de Córdoba | ARG | European J of Plant Path | 1 | 1.0 |
| Changes of lipids composition in different ontogenetic stages of *Thecaphora frezii*: expression of key enzymes for lipid biosynthetic pathways | 2022 | Diaz | CEPROCOR | ARG | Journal of Applied Microbiology | 0 | 0 |
| First draft genome of *Thecaphora frezii*, causal agent of peanut smut disease | 2023 | Arias | USDA | US | BCM Genomic Data | 0 | 0 |

